# PIWIL1 is recruited to Centrosomes during Mitosis in Colorectal Cancer Cells and is linked to cell cycle progression

**DOI:** 10.1101/2024.06.04.597425

**Authors:** M.R. Garcia-Silva, S. Montenegro, S. Dacosta, J.P. Tosar, A Cayota

## Abstract

PIWI proteins, traditionally associated with germline development, have recently gained attention for their expression in various cancers, including colorectal cancer (CRC). However, the molecular mechanisms underlying their reactivation and impact on cancer initiation and progression remain elusive. Here, we found that PIWIL1 is expressed at relatively high levels in CRC-derived samples and cell lines, where it undergoes a dynamic re-localization to the centrosome during mitosis. Knockdown of PIWIL1 induces G2/M arrest associated to disruption of mitotic spindle and aberrant metaphase events, highlighting its role in cell cycle progression. We have also found that expression of PIWIL1 is lost during differentiation of Caco-2 cells into enterocytes and that PIWIL1 is expressed in cells at the base of intestinal crypts in normal human colon tissue, where intestinal stem cells are known to reside. Thus, it is possible that the presence of PIWIL1 in cancer cells reflects a physiological role of this protein in stem cell maintenance, what would argue in favor of the proposed stem cell origin of CRC. Supporting this view, dedifferentiation of human fibroblasts into induced pluripotent stem cells (iPSc) implies reactivation of PIWIL2 expression, another member of the PIWI protein family. Overall, our findings suggest a role of PIWIL1 in mediating cell cycle dynamics, both in colorectal cancer cells and possibly also in intestinal stem cells. In a broader aspect, we provide support for an involvement of PIWI proteins in somatic stem cell maintenance, expanding nongonadal functions for this protein family.

## Introduction

Colorectal cancer (CRC), one of the leading causes of cancer-related mortality worldwide, has been extensively studied for its intricate underlying molecular bases. Despite this, CRC remains a global health challenge, demanding deeper exploration of the particular molecular pathways driving its progression and the identification of novel therapeutic targets (Patel et al., 2022). Among the myriad pathways implicated in cancer progression, the pathway mediated by PIWI (P-element-induced wimpy testis) proteins, traditionally associated with germline development and maintenance of genome stability, has emerged as a captivating focus of investigation due to its overexpression in various cancers, including CRC (Wang et al., 2022).

Notably, PIWIL1, a key component of piRNA-mediated gene expression regulation pathways, has gathered attention for its suspected role as an oncogene since 2002, when it was first reported to be overexpressed in seminoma (a testicular-derived tumor) (Qiao et al., 2002). Nowadays, overexpression of different PIWI proteins has been frequently observed in a variety of cancers, including gastric cancer, sarcoma, esophageal squamous cell carcinoma, endometrial cancer, colon cancer, cervical cancer, glioma, hepatocellular carcinoma, ovarian cancer, breast cancer, bladder cancer, and renal cell carcinoma (Chakravorty, 2022). Nevertheless, the mechanisms underlying PIWI protein overexpression and their impact on the initiation and progression of cancer remain elusive.

Reports on the expression of piRNAs in either somatic or neoplastic cells are at times conflicting. In this respect, we have recently reported that a large fraction of the so-called somatic piRNAs, which have been described in nongonadal tissues, derive from non-coding RNAs and do not bear the specific molecular properties of canonical piRNAs (Tosar et al., 2018, 2021). Additionally, experimental data obtained in CRC cells by two independent groups could not find evidence for the expression of canonical piRNAs linked to PIWIL1 (Genzor et al., 2019 and Li et al., 2020). This suggests that PIWI proteins could act independently of piRNAs in these contexts. However, how piRNA-independent molecular mechanisms of PIWI proteins contribute to cancer initiation and progression is not clear yet.

Henceforth, we aimed to shed light on the non-canonical functions of PIWIL1 re-activation in CRC cancer models. Our analysis first showed an overexpression of *PIWIL1* in CRC patient’s samples from TGCA database. In addition, subcellular localization of PIWIL1 unveiled its remarkable re-localization during the cell cycle, from a nuclear scattered/dispersed granular distribution in interphase, to a microtubule-organizing center (MTOC)-like localization during mitosis in human cell lines derived from colorectal cancer (Caco-2). Building upon this observation, we delved into the functional consequences of PIWIL1’s presence at the centrosome during mitosis. Our data clearly demonstrate that PIWIL1 silencing leads to a distinct cellular phenotype, characterized by cell cycle stalling at G2/M phase and the accumulation of aberrant metaphase events in Caco-2 cells. Remarkably, in normal colon-derived tissue samples, PIWIL1 is observed at the base of colon crypts, a known niche of intestinal stem cells. This is consistent with the PIWI pathway being relevant for the maintenance of stemness and self-renewal in somatic stem cells (Palakodeti et al., 2008; Juliano et al., 2014; Galton et al., 2022). Also supporting a role in stemness maintenance, PIWIL1 expression seems to be downregulated during differentiation of Caco-2 cells to enterocytes. Albeit for a different PIWI protein (PIWIL2), we also observed dedifferentiation-induced expression in a model of induced pluripotent stem cell (iPS) generation, confirming our hypothesis that PIWI proteins are important for human stem cell maintenance.

Mechanistically, our findings support an association between PIWIL1 and the mitotic spindle, as previously observed in germ cells (Rodriguez et al., 2005; Venkei et al., 2020). Taken together, our results highlight the relevance of PIWI proteins in the regulation of the cell cycle in undifferentiated cells, possibly explaining their unexpected but widely documented expression in cancer.

## Results

### PIWIL1 is Upregulated in primary tumor-derived samples from Colorectal Cancer Patients Compared to Healthy Individuals

Data mining was performed to determine the expression of the four PIWI proteins codified by the human genome, in samples-derived from patients with CRC. *PIWIL1, PIWIL2, PIWIL3* and *PIWIL4* RNA-seq data from colon primary tumor and normal cohorts-derived samples were analyzed using the UCSC Xena Browser (Goldman et al., 2020), integrating data from TCGA, TARGET and GTEX databases. RNA-seq data from 596 samples were ranked according to *PIWI* gene expression levels, revealing a positive correlation between *PIWIL1* and tumor status (Figure 1A-C).

**Figure 1:**
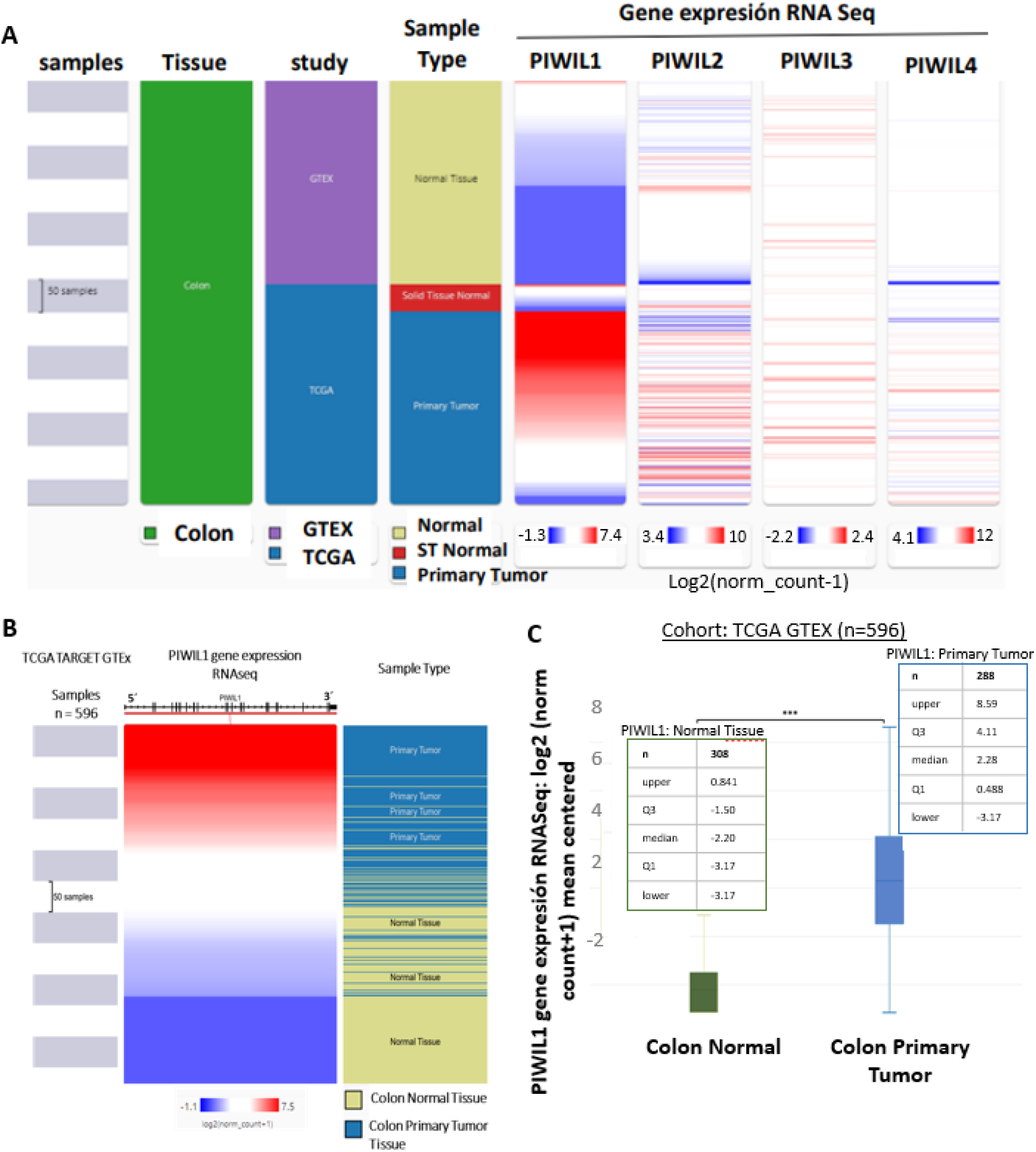
PIWIL1 is overexpressed in CRC tissue samples. **A**. Visual Spreadsheet obtained from https://ucscxena.github.io for PIWIL1-4, in normal colon tissue (GTEX) and TCGA (CRC samples). N = 650 samples. **B**. UCSC Xena Visual Spreadsheet for *PIWIL1* mRNA in normal and primary tumor-derived samples. **C**. *PIWIL1* levels in cancer tissue samples from TCGA and normal colon tissue from GTEx (p ≤ 0.0001).

### PIWIL1 is localized in the nucleus in colon cancer cells

Cell lines derived from human colon carcinoma and the beningn colon epithelial cell line (CCDCoN) were assessed for PIWIL1 expression at transcript and protein level. All cell lines expressed variable levels of PIWIL1 transcripts, measured by RT-PCR, and protein levels assesed by immunoprecipitation followed by Western blot (Figures 2A and 2B, respectively). Indirect immunofluorescence (IF) to determine PIWIL1 subcellular localization showed a predominant nuclear localization in Caco-2 (Figure 2C) and other CRC cell lines (Figure 2D), with the signal presenting a patterned distribution in the nucleoplasm and a remarkable association with the nuclear lamina during interphase.

**Figure 2:**
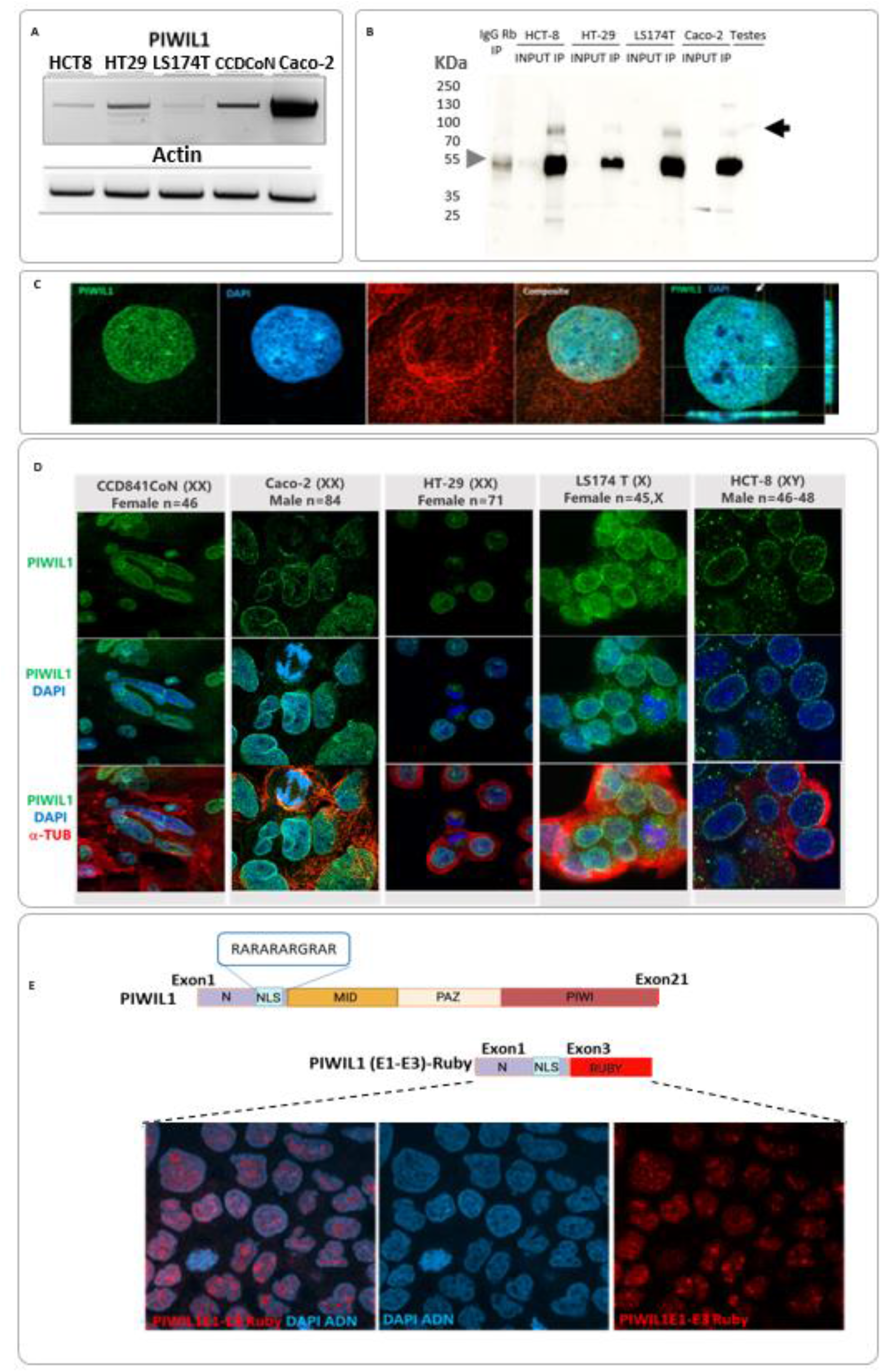
PIWIL1 is expressed in several CRC cell lines and localizes in the nucleus. **A**. RT-qPCR of full-length (2500bp; Genbank Gene ID:9271) *PIWIL1* in different CRC cell lines and actin as loading control. **B**. Western blot of immunoprecipitated fractions enriched in PIWIL1 from different CRC cell lines. Positive control: testicular tissue. Black arrows: PIWIL1 (100 KDa). Gray arrows: IgG. **C**. PIWIL1 localization in interphasic Caco-2 cells, measured by indirect immunofluorescence (green). Blue: DAPI. Red: Beta tubulin. Last panel: orthogonal sectioning of a confocal z-stack image confirming intranuclear PIWIL1 expression. **D**. Representative images in different CRC-derived cell lines and normal colon-derived cell line (CCD841CoN). **E**. Diagram of the construction performed to test PIWIL1 predicted NLS (blue box) tagged to mRuby (top) and widefield microscopy showing nuclear expression of PIWIL1 NLS-tagged mRuby (red) in interphasic Caco-2 cells (bottom)

*Drosphila* Piwi shuttles to the nucleous thanks to a N-terminal nuclear localization signal (NLS) that is exposed upon piRNA binding (Yashiro et al., 2018). Interestingly, we identified a predicted NLS in the exon 2 of PIWIL1 (aa 4 to 14), comprising a stretch of basic residues (arginine and alanine). To determine its functionality as a *bona fide* NLS in Caco-2 cells, we fused this sequence motif to mRuby2. Consistent with the observed predominantly nuclear expression of PIWIL1 in these cells (Figure 2C), NLS-tagged mRuby2 was mostly expressed in the nucleus (Figure 2E).

### PIWIL1 localizes to the centrosome in Caco-2 cells during mitosis

While PIWIL1 was mostly localized in the nucleoplasm or in the nuclear lamina in interphasic cells, we consistently observed its expression in the centrosome of mitotic cells (see, for example, Figure 3). With the purpose of better elucidating the subcellular distribution of PIWIL1 during the cell cycle, we synchronized Caco-2 cells and confirmed PIWIL1 is dynamically redistributed during mitosis (Figure 3A). Confirming previous observations, synchronized Caco-2 cells co-stained for beta tubulin showed PIWIL1 relocalization to the MTOC during mitosis (Figure 3B).

**Figure 3:**
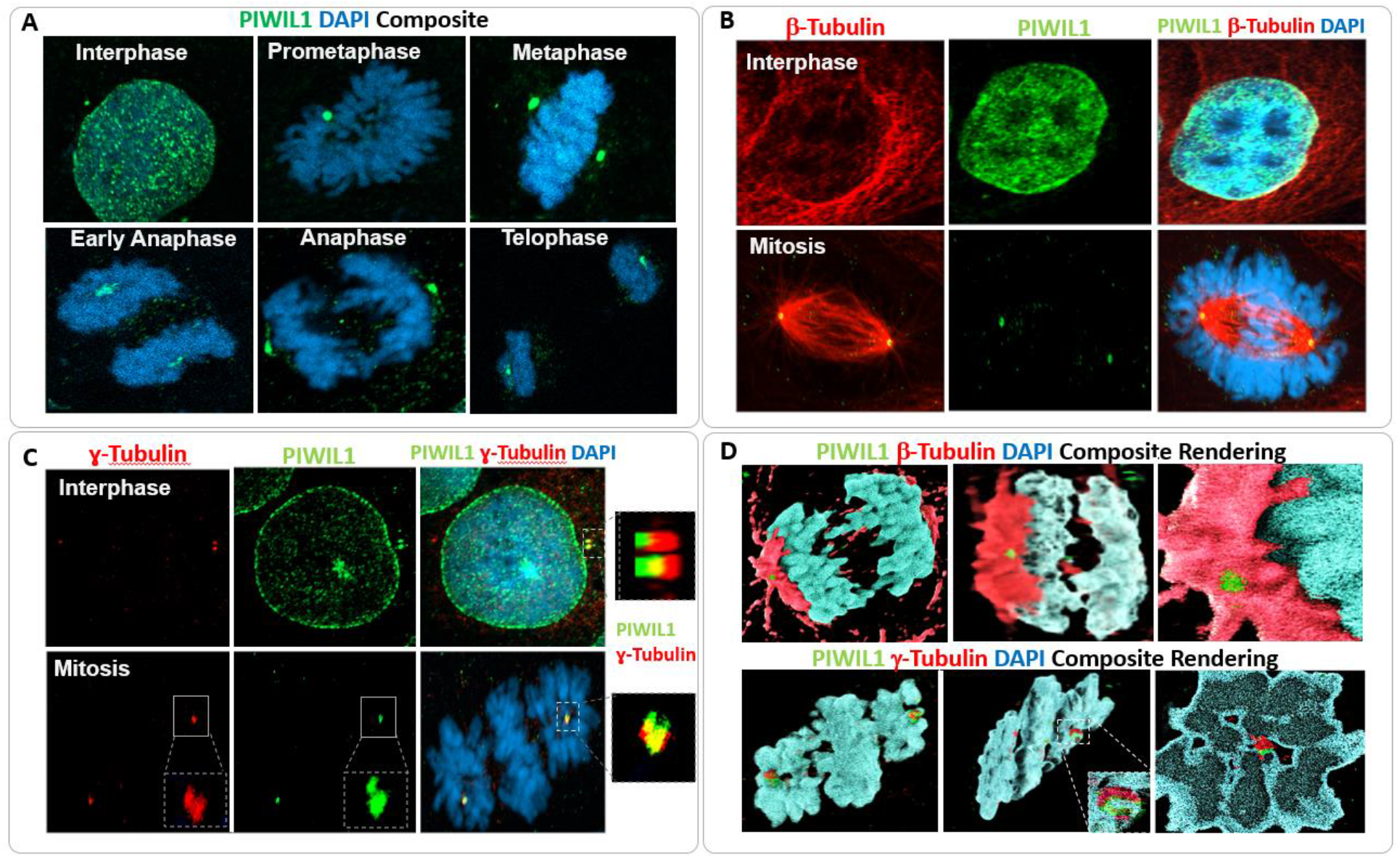
PIWIL1 is recruited to centrosomes in mitotic Caco-2 cells. **A**. PIWIL1 localization in synchronous cultures of Caco-2 cells (green). Images representing different stages of mitosis are shown, based on DNA staining with DAPI (blue). **B**. Subcellular localization of PIWIL1 (green) and beta tubulin (red) in two representative images of Caco-2 cells in interphase or mitosis. Note the aberrant mitotic spindle that is characteristic of this cell line. **C**. Subcellular localization of PIWIL1 (green) and the centrosome marker gamma tubulin (red), in interphase and mitosis. Inset: magnified view of the centrosome. **D**. Rendering images (made in AGAVE) evidencing PIWIL1 recruitment to the MTOC (upper panel) and co-localization with centrosomes (bottom panel).

Co-localization assays of PIWIL1 and gamma tubulin, a known centrosomal marker, confirmed PIWIL1 as an integral component of the MTOC or the centrosome (Figure 3C). 3D Rendering of composite images are depicted in Figure 3D, where PIWIL1 and gamma tubulin appear to be arranged in a manner reminiscent of two interlocking rings, fitting together seamlessly.

### PIWIL1 silencing causes G2/M arrest and aberrant mitosis

To understand the functional implications of PIWIL1 localization to centrosomes in dividing CRC cells, we performed siRNA-mediated PIWIL1 knock down (KD). Levels of PIWIL1 protein and mRNA decreased by 70% as assessed by fluorescence microscopy and RT-qPCR, respectively (Figure 4A). Surprisingly, KD cells showed a significantly higher frequency of mitotic defects (vs control siRNA-transfected cells), comprising multiple centromeres, anormal mitotic spindles or misaligned metaphasic chromosomes (Figure 4B). This was followed by cell cycle arrest at G2/M, as measured by flow cytometry using propidium iodide DNA staining (Figure 4C). These cell division defects were consistent with KD cells having a lower proliferation rate and / or migratory capacity, as measured by wound healing assays (Figure 4D). In sum, despite Caco-2 cells are transformed and polyploid, they seem to rely on PIWIL1 to regulate mitotic fidelity, potentially avoiding mitotic catastrophe.

**Figure 4:**
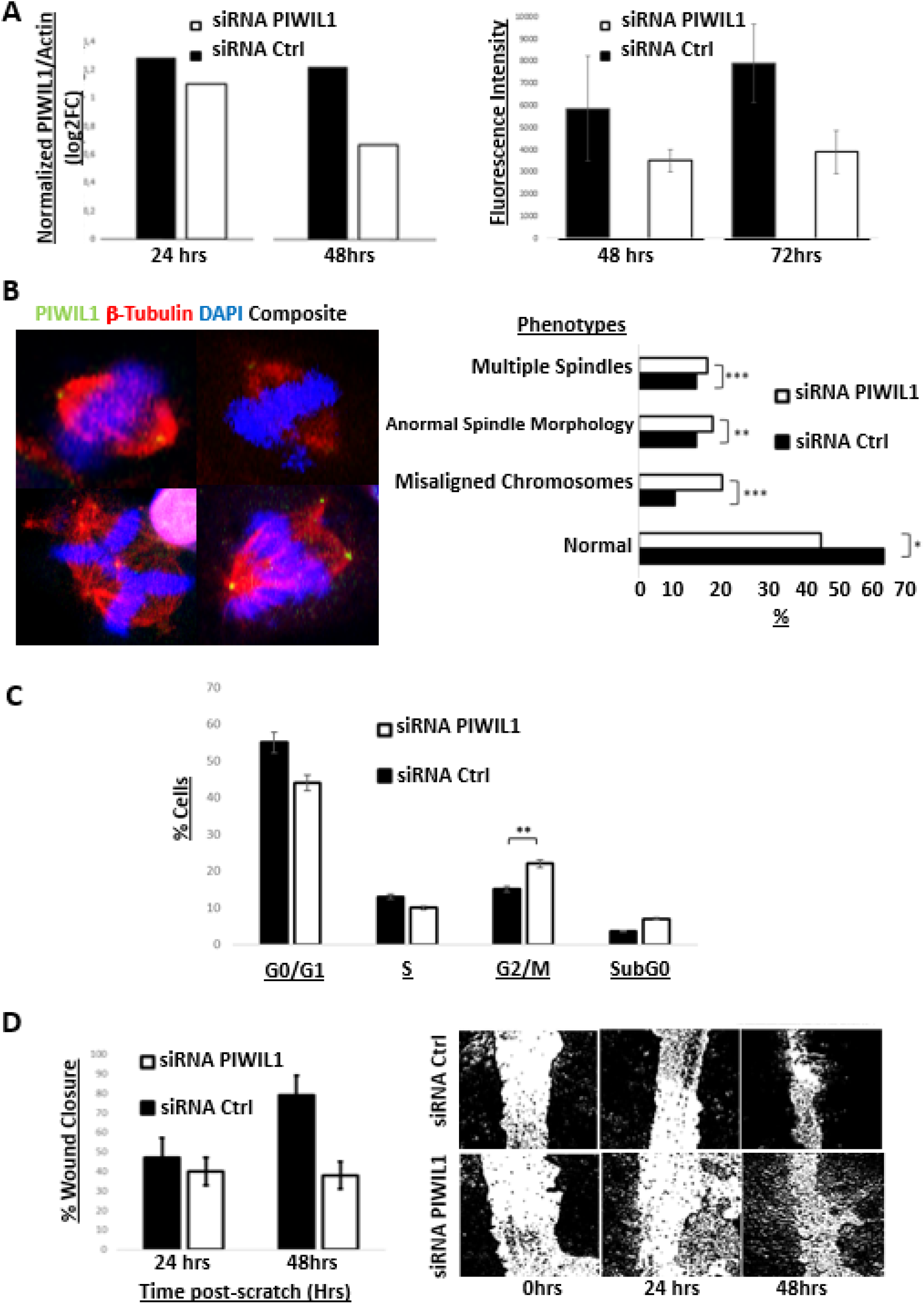
PIWIL1 silencing leads to aberrant mitosis and G2/M arrest. **A**. Relative *PIWIL1* mRNA levels (normalized to actin; left panel) and protein levels, quantified based on fluorescence intensity (right panel), in PIWIL1 KD cells vs cells transfected with a control siRNA. **B**. Left panel: representative images of mitotic Caco-2 cells after PIWIL1 knock down. Tubulin (Red) and DAPI (Blue) are labeled to quantify mitotic abnormalities. Right Panel: Percentage of observed phenotypes. N= 100 cells in mitosis for each condition (PIWIL1 KD vs control siRNA). **C**. Cell cycle assays assessed by flow cytometry (PI staining to quantify DNA content). **D**. Wound-Healing assay comparing the capacity of PIWL1 KD Caco-2 cells vs control siRNA-transfected cells to close a scratch at t = 24 or t = 48 h. Representative images are shown in the right. N = 3 for each condition.

### PIWIL1 expression is restricted to stem cell niches in normal human colon tissue and is lost upon differentiation

Next, we asked whether PIWIL1 expression in CRC patients and CRC cell lines is due to an aberrant expression of this otherwise germline-specific protein, or whether PIWIL1 is expressed in a subset of human colon cells under physiological conditions. Surprisingly, immunohistochemistry and immunofluorescence microscopy of normal human colon tissue showed PIWIL1 staining restricted to cells at the bottom of each intestinal crypt, where stem cells are known to be located (Ritsma et *al*, 2014) (Figure 5A). In contrast, colon cancer tissue showed pervasive PIWIL1 staining, with virtually all cells staining positive. However, unlike CRC cell lines, the signal was predominantly cytoplasmic in both healthy and cancer tissues.

**Figure 5:**
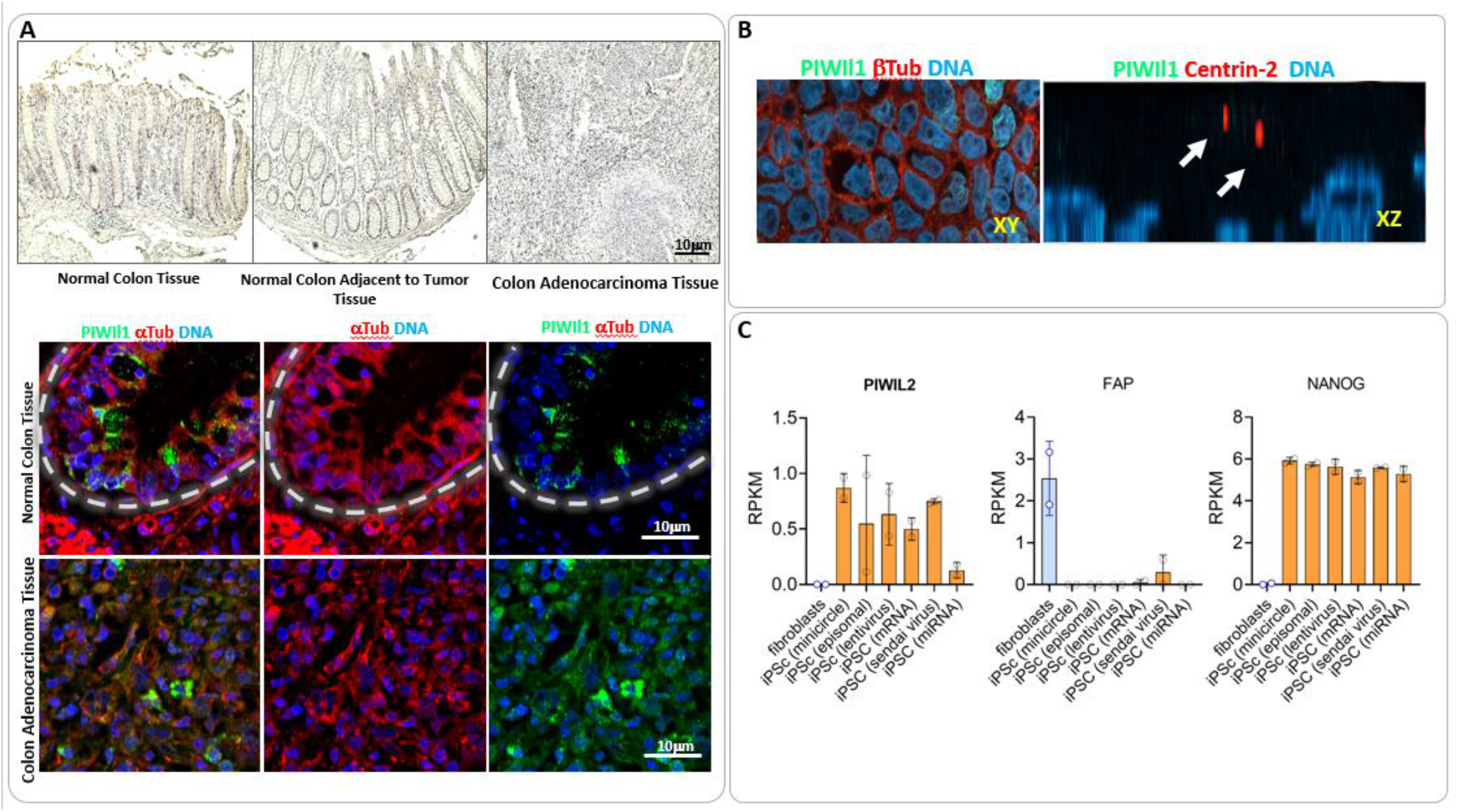
PIWIL1 expression anti-correlates with the differentiation status of colon epithelial cells. **A**. Upper panel: Immunohistochemistry staining of PIWIL1 in tissue sections from normal colon tissue (left), peritumoral normal colon tissue (center) and colon adenocarcinoma (right). Low levels of Hematoxiline and Eosine were used to stain cells and nucleus. Bottom panel: Immunofluorescence of PIWIL1 (green) in the stem cell-containing basal region of normal colon crypts (top) and colon adenocarcinoma-derived tissue (bottom). Composites images for PIWIL1 (green), DAPI (blue) and tubulin (red) are shown. White dashed lines demarcate the limits of the intestinal crypts. **B**. Immunofluorescence of PIWIL1 in Caco-2 cells submitted to differentiation in transwell assays. PIWIL1 is absent when we observed polarized, differentiated enterocytes, characterized by centrin-2 labelling of apical centrosomes. **C**. Analysis of RNA-seq data from GEO: GSE69626, showing *PIWIL2, FAP* and *NANOG* mRNA abundance (expressed as Reads per Kilobase per Million mapped reads; RPKM) in differentiated human fibroblasts and induced pluripotent stem cells (iPSc) obtained from those fibroblasts using different methods.

Intestinal crypts, which form the base of the villi, are the basic structural and functional units of intestinal epithelium. At the bottom of each crypt, a small number of about eight stem cells (Ritsma et al, 2014) are constantly dividing to give place to transit-amplifying progenitor cells. These progenitor cells divide as they migrate upwards and differentiate into two distinct lineages: enterocytes (absorptive cells) and secretory cells.

Having observed that, in normal colon tissue, PIWIL1 expression is restricted to the stem cell population residing at the base of intestinal crypts, we asked whether PIWIL1 expression is downregulated during progenitor cell differentiation into enterocytes. To study this, we followed a well-stablished protocol (Hubatsch et al., 2007) where Caco-2 cells are differentiated to enterocytes by means of a transwell assay. Surprisingly, unlike 2D cultures of undifferentiated cells (Figure 2), we were not able to find PIWIL1 expression when Caco-2 cells were differentiated to enterocytes (Figure 5B). To validate the differentiation protocol, we analyzed the subcellular localization of centrin-2, a structural component of the centrosome, that migrates to the apical level of the cells upon differentiation (Figure 5B). Thus, loss of PIWIL1 during Caco-2 differentiation resembles the situation in normal colon epithelia, where only undifferentiated cells residing at the base of intestinal crypts express this protein to detectable levels.

To study whether PIWI proteins are required for the maintenance of stemness and self-renewal in non-transformed cells, we analyzed RNA-seq data from GEO: GSE69626. In this study, human fibroblasts were de-differentiated into induced pluripotent stem cells (iPSCs) using six different reprogramming methods (Churko et al. Nature Biomed. Eng. 2017). Surprisingly, while *PIWIL1* was not involved in this process, *PIWIL2* expression was induced in all reprogrammed cells (Figure 5C). Differentiated fibroblasts expressed *FAP* but lacked *PIWIL2* and the stem cell marker *NANOG*. Conversely, de-differentiated cells expressed *PIWIL2* and *NANOG*, while silencing *FAP*.

Albeit involving a different PIWI protein to the one studied herein, results above strengthen the link between PIWI protein expression and stem cell biology. This is in agreement with PIWIL1 expression in stem cells from healthy colon tissue. In addition, these results potentially explain PIWIL1 overexpression in CRC based on the potential stem cell origin of colorectal cancer (Huels and Sansom et al. British J of Cancer, 2015).

## Discussion

Several breakthroughs in the understanding of the PIWI pathway beyond its traditional involvement in transposon signaling in the gonads have been made in the past decade (Ross et al., 2014; Dai et al., 2019; Ramat et al., 2020). However, a complete understanding of extragonadal PIWI protein function, particularly in cancer, remains incompletely understood. In this regard, reactivation of PIWI expression has been attributed to aberrant DNA methylation, a hallmark of various types of tumors (Litwin et al., 2017). Aberrant expression of other genes from the PIWI pathway has been observed in several cancer cell lines, in the absence of a concomitant reactivation of piRNA biogenesis (Genzor et al. PNAS 2019).

Our own data-mining analysis of 600 patients’ samples obtained from TGCA and GTEX showed pervasive PIWIL1 expression in CRC compared to normal colorectal tissue. Our results are therefore in good agreement with previous studies reporting aberrant PIWIL1 expression in CRC patients (Liu et al., 2012; Sun et al., 2017; Raeisossadati et al., 2014). Interestingly, a piRNA-independent function for PIWIL1 in cancer cell proliferation has been previously reported in human pancreatic cancer cell lines, where it functions as an activator of the E3 ubiquitin ligase anaphase promoting complex/cyclosome (Li et al., 2020). In addition, PIWIL1 overexpression within an adenovirus vector promoted cell proliferation, associated with changes in global DNA methylation levels (Yang et al., 2015). However, the molecular mechanisms linking PIWIL1 expression and sustained proliferation have not been thoroughly explored.

In our study, we observed physical association between PIWIL1 and centrosomes during mitosis in Caco-2 cells and other CRC cell lines. PIWIL1 silencing in these cells induced cell cycle arrest, diminished cell proliferation, and provoked the accumulation of mitotic defects. These observations are in good agreement with earlier studies linking PIWIL1 with the anaphase-promoting complex, a key player in cell cycle control (Li et al., 2020). These findings suggest that cell cycle regulation mediated by PIWIL1 might be mediated through interactions with other proteins, regardless of piRNAs. Interestingly, during murine male meiosis, PIWIL1 is involved in the control of meiotic segregation and its deficiency causes chromosome misalignment and mis-segregation, leading to aneuploidy via its contact with transcribed centromeric regions (Hsieh et al., 2020). Our findings of PIWIL1 recruitment to the centromere/MTOC during mitosis in CRC cells, and its importance for proper chromosome alignment during metaphase, support the idea that at least some of the gonadal functions of PIWIL1 are conserved in somatic tissues. These insights underscore the multifaceted role of PIWIL1 in cell cycle regulation and support previously recognized piRNA-independent functions during cell division.

Our data also agrees with previous work conducted in gastric cancer cells. For instance, Liu et al. found that PIWIL1 downregulation effectively inhibited proliferation and induced G2/M phase cell cycle arrest (Liu et al., 2006). Additionally, in breast cancer and sarcoma cell lines, PIWIL1 overexpression increases cyclin levels and regulates cyclin-dependent kinase inhibitors p21 and p27 (Siddiqi et al., 2012; Cao et al., 2016). In glioblastoma, PIWIL1 knockdown induced significant changes in cell cycle-related genes, confirming the dysregulation of cyclin levels and the increase in cells arrested in G2/M (Huang et al., 2021).

The G2/M cell cycle transition slowdown observed as a result of reduced PIWIL1 expression in Caco-2 cells could have several functional implications. One hypothesis is that loss of PIWIL1 leads to a reduced proliferative capacity as a response to increased genomic instability and/or mitotic spindle disruption. Another hypothesis highlights the role of PIWI proteins in cancer development and progression by promoting a stem cell-like status that sustains self-renewal and proliferation (Litwin et al., 2017). In the context of CRC, the origin of the malignancy is not well understood, but it is likely that the cells that give rise to the tumor are actually intestinal stem cells (Huels and Sansom, 2015). Sousa-Victor P. *et al* have identified PIWI as a critical regulator of somatic stem cell function in intestinal stem cells in flies, safeguarding maintenance of stemness (Sousa-Victor et al., 2017). In humans, others have observed a positive correlation between PIWI pathway genes and cancer stem cell markers in CRC (Litwin et al., 2015).

PIWIL1 localization within colon stem cell niches in healthy intestinal tissue, and its overexpression in colon cancer cells, might reflect an evolutionary path leading from colon stem cells to colon cancer. In fact, the ability to *in vitro* differentiate Caco-2 cells into enterocytes could be considered evidence that these are non-terminally differentiated cells. Consistent with a role restricted to undifferentiated or stem cells, PIWIL1 was lost during Caco-2 differentiation to enterocytes. Taken together, these data suggest that the presence and activity of PIWIL1 at the base of intestinal crypts in normal colon tissue is being reflected in CRC-derived cells, either due to the stem cell origin of the CRC cell lines used herein, or due to functional convergence in the adoption of stem cell-like characteristics and undifferentiated states.

The expression of PIWIL1 in colon tumor cells seems to be a necessary event for proper cell division. Its absence leads to aberrant mitosis and increases genomic instability, as demonstrated by previous studies and our own findings. Our study thus unveils an opportunity for targeting PIWIL1 as a novel CRC therapeutic strategy, potentially leading to mitotic catastrophe in these cells.

In conclusion, PIWIL1 might be important for stem cell maintenance in the human intestine. For the same reason it might also be frequently expressed in CRC, where it seems to play a role in mitotic fidelity and/or cell cycle regulation. This protein seems to be silenced during differentiation, while its downregulation induces genomic instability in cancer cells. Therefore, we would like to make a case for exploring PIWIL1 as an attractive therapeutic target in the future.

## Materials and Methods

### Bioinformatic Resources

Bioinformatic data mining was performed in CRC cohorts in TCGA, GTEX and TARGET in order to asses PIWIL1 gene expression. We used Xena’s Visual Spreadsheet and Chart View to construct Figure 1 A and B to analyze PIWIL1 upregulation in connection with CRC (Goldman et al., 2020).

### Cell Culture and qRT-PCR

Caco-2 (HB-37) cells were purchased from the American Type Culture Collection (ATCC) and cultured in DMEM (Gibco) with 10% fetal bovine serum (Gibco) according to the supplier’s instructions. Caco-2 cells were synchronized with two pulses of 2mM of Thymidine (Sigma) for 16 hours.

Total RNA was isolated using TRIzol reagent (Invitrogen). cDNA was obtained by MLV Reverse Transcriptase (Merck) using 1µg of Total RNA and an oligo (dT)_18_ primer. The primer sequences for PIWIL1 RT–qPCR are: ATGGCGTACAGACACGAGGC (Forward) and AGTGACAGATTTGGCTCTCTG (reverse).

### Immunofluorescence and Confocal Microscopy

Caco-2 cells (1×10^6^) were plated onto microscope slides in a 24-well plate. After the desired time, cells were fixed using 4% paraformaldehyde for 15 min at RT and permeabilized with PBS-tween 0.2% for 3 minutes. Then, cells were incubated with blocking buffer (5% BSA in PBS) for 1 hour at 37°C. The cells were washed three times with PBS and incubated with primary antibody diluted in blocking buffer overnight in a humid chamber. Fluorophore-conjugated secondary antibodies were incubated, after 3 washes with PBS, on the slides 1 hour. DNA staining was performed with 300 µM DAPI (Sigma) for 5 minutes at room temperature in the dark. After washing with PBS, coverslips were mounted using Prolong Gold Antifade Reagent (Thermo Fisher Scientific). Cells were photographed with a Zeiss confocal microscope LSM 880 and analyzed using Fiji-ImageJ software. Rendering of images obtained by confocal Z Stacks was performed in AGAVE path-trace viewer (Allen Institut for Cell Science).

Primary antibodies used in this study were: anti-PIWIL1 HPA018798 (Sigma-Prestige) and HPA018798 (Abcam). Anti β−tubulin sc-55529 (Santa Cruz tech). γ−Tubulin (SIGMA T6557). Centrin-2 (Merck, clone 20H5).

### PIWIL1 knock down assays and cell cycle flow cytometry

Caco-2 cells were transfected with siRNA (10nM) to PIWIL1 (Trifecta Kit; IDT), using Lipofectamine 3000 (Invitrogen) according to manufacturer’s instructions. The mRNA expression levels of PIWIL1 and Actin were examined by semiquantitative RT-PCR from total RNA using TriZol Reagent (Sigma) 72 h after siRNA transfection.

Cell cycle progression of Caco-2 cells was measure 72 h after siRNA PIWIL1 *vs* irrelevant scrambled siRNA transfection by flow cytometry with Propidium Iodide (PI) Staining. Briefly, 1 million Caco-2 cells were fixed with cold 70% ethanol for 30 min, washed twice in PBS, and treated with Ribonuclease I (100μg/mL Sigma). After RNase treatment, cells were stained with 50μg/mL PI (Invitrogen), and analyzed using an AttuneTM NxT instrument and the FlowJo software. Approximately 10,000 cells per condition were selected after the gating strategy. The first gate was placed on FSC vs SSC. The second gate was placed on PE vs SSC-A and displayed as an histogram using PE intensity. Percentage of cells in each cell cycle phase was calculated using markers set within the analysis program.

### Wound Healing Assays

1×10^4^ Caco-2 and PIWIL1 knockdown Caco-2 cells were grown in 24-well plates. After the cells have become confluent we performed a cell-free gap in the monolayer by mechanical scratching (scratch wound) with a white sterile pipet tip. optical microscopy can be used to observe cells migrating into the wound area. Imaging Data was acquired every 24 hours using phase-contrast imaging of the wound area in the field-of-view using 10X objective lens. % of wound closure (Area) was measure by the wound healing tool by Volker Baecker for ImageJ.

### Colon-derived Tissue Arrays Immunohistochemistry

Colon carcinoma tissue array with matched adjacent normal colon tissue as control, including TNM, clinical stage and pathology grade, was acquired from Tissuearray.com (CO243b). Before immunohistochemistry, antigen retrieval was performed baking slides for at least 60 minutes at 60°C before de-paraffinization using xylene. De-paraffinized slides were treated as described for the immunofluorescence protocol.

### Caco-2 differentiation using Transwell Experiments

1.0×10^5^ Caco-2 cells were seeded on HTS-24 Transwell-Plates containing a 0.4 μm membrane (Costar; Corning Inc). Experiments were performed in triplicates. Plates were incubated at 37°C overnight and the medium of the upper chamber was replaced by new DMEM-PEST to remove non-adherent cells and reduce the risk of multilayer formation. The culture medium was refreshed every second day in both compartments for a total of 21 days. Polarized monolayer cells were checked by actin labelling and the formation of vertical filaments. Membrane filters were disassembled and proceed as detailed in the immunofluorescence protocol.

## Acknowledgements

Authors gratefully acknowledge the Advanced Bioimaging Unit and the Cell Biology Unit (Cytometry platform: MSc. Paula Céspedes) at Institut Pasteur of Montevideo for the support and assistance in the present work.

This work was supported by FOCEM (Fondo para la Convergencia Estructural del Mercosur) (COF 03/11), ANII (PEC_3_2019_1_158811) for the AttuneTM NxT equipment and to Comisión Sectorial de Investigación Científica (CSIC) Programa Grupos 2022 (Área salud).

